# Structural basis for bispecific antibody design: arrangement of domain linkage produces activity enhancement

**DOI:** 10.1101/2024.04.25.591206

**Authors:** Kyohei Sato, Shiro Uehara, Atsushi Tsugita, Shieru Ishiyama, Atsushi Maejima, Ishin Nakahara, Misae Nazuka, Takashi Matsui, Christos Gatsogiannis, Takeshi Yokoyama, Izumi Kumagai, Koki Makabe, Ryutaro Asano, Yoshikazu Tanaka

**Affiliations:** Graduate School of Life Science, Tohoku University; Graduate School of Science and Engineering, Yamagata University; Graduate School of Engineering, Tokyo University of Agriculture and Technology; School of Science, Kitasato University; Center for Soft Nanoscience, Institute for Medical Physics and Biophysics, University of Münster, 48149 Münster, Germany; The Advanced Center for Innovations in Next-Generation Medicine (INGEM), Tohoku University

## Abstract

A bispecific antibody (BsAb) is a protein genetically engineered from two different antibodies, allowing simultaneous binding to two kinds of antigen to bring them into close proximity. BsAbs have been developed as anti-cancer drugs that accumulate lymphocytes onto cancer cells by bridging antigens present on each. Ex3 is a bispecific diabody composed of the two fused variable regions (Fvs) of an anti-epidermal growth factor receptor (EGFR) antibody and an anti-CD3 antibody with potent cancer cytotoxic activity. In Ex3, the LH-type, in which the variable regions of the light chain (VLs) are located at the N-terminus of those of the heavy chain (VHs), exerted 1000-fold greater anticancer activity than the HL-type, in which the VHs are located at the N-terminus of the VLs. This effect (termed ‘activity enhancement’), in which the activity is greatly enhanced by domain rearrangement, has been reported not only for Ex3 but also for several other BsAbs. However, the molecular details of this activity enhancement have yet to be elucidated. In this study, we determined the cryo-EM structures of Ex3 LH- and HL-types in complex with CD3 and EGFR. Structural comparison of the LH- and HL-types showed that rearrangement of the domain linkage produces drastic structural differences in the overall shape of these complexes, and dynamics attributed to the flexibility between the two Fvs. These findings provide valuable insights into the molecular mechanism for the activity enhancement of BsAbs. This study will be a stepping stone towards establishing a design foundation for BsAb development.

## Introduction

Due to their high specificity to targets, the design of antibodies is valuable for drug development and forms one of the main modalities for cancer therapy^1^. In addition, various types of recombinant antibodies have been actively developed in recent years to create next-generation antibodies with higher functionality^2,3,4^. A bispecific antibody (BsAb) is a recombinant antibody genetically engineered by fusing the variable regions of two different antibodies, which consequently can bind to two different antigens simultaneously. To date, several effective anti-cancer BsAbs have been reported^5,6^.

Ex3^7^ is a bispecific diabody composed of the variable regions (Fvs) of anti-epidermal growth factor receptor (EGFR) antibody 528^8,9^ and an anti-CD3 antibody OKT3^10^. 528 binds to EGFR on the surface of cancer cells, while OKT3 binds to CD3, an accessory molecule of the T cell receptor on the surface of T cells. Therefore, Ex3 can form a bridge between cancer cells and T cells, inducing the accumulation of activated T cells on cancer cells and consequently exhibit cancer cytotoxicity. In the design of BsAbs, the order and orientation of the domain linkages can be changed. Several domain configurations exist even in the same combination of BsAb. In Ex3, the LH-type, in which the variable regions of the light chain (VLs) are located at the N-terminus of the variable regions of the heavy chain (VHs), has been shown to have 1000-fold greater anticancer activity than the HL-type, in which VHs are located at the N-terminus of VLs^11^ (Fig. 1a). However, the binding affinity and bridging ability to both soluble antigens is, interestingly, comparable between HL- and LH-types^11^. Also, it has been reported that the ability to bridge cancer cells and T-cells is enhanced in the LH-type^12^, which correlates with its cytotoxic activity. These observations suggest that the difference in activity is due to the dynamics of the cell surface rather than the antibody-antigen affinity itself. A similar phenomenon, in which the activity is greatly enhanced by domain rearrangement (termed ‘activity enhancement’), has been reported for several other BsAbs in addition to Ex3^13,14^ However, the detailed mechanism has not been elucidated.

**Figure 1:**
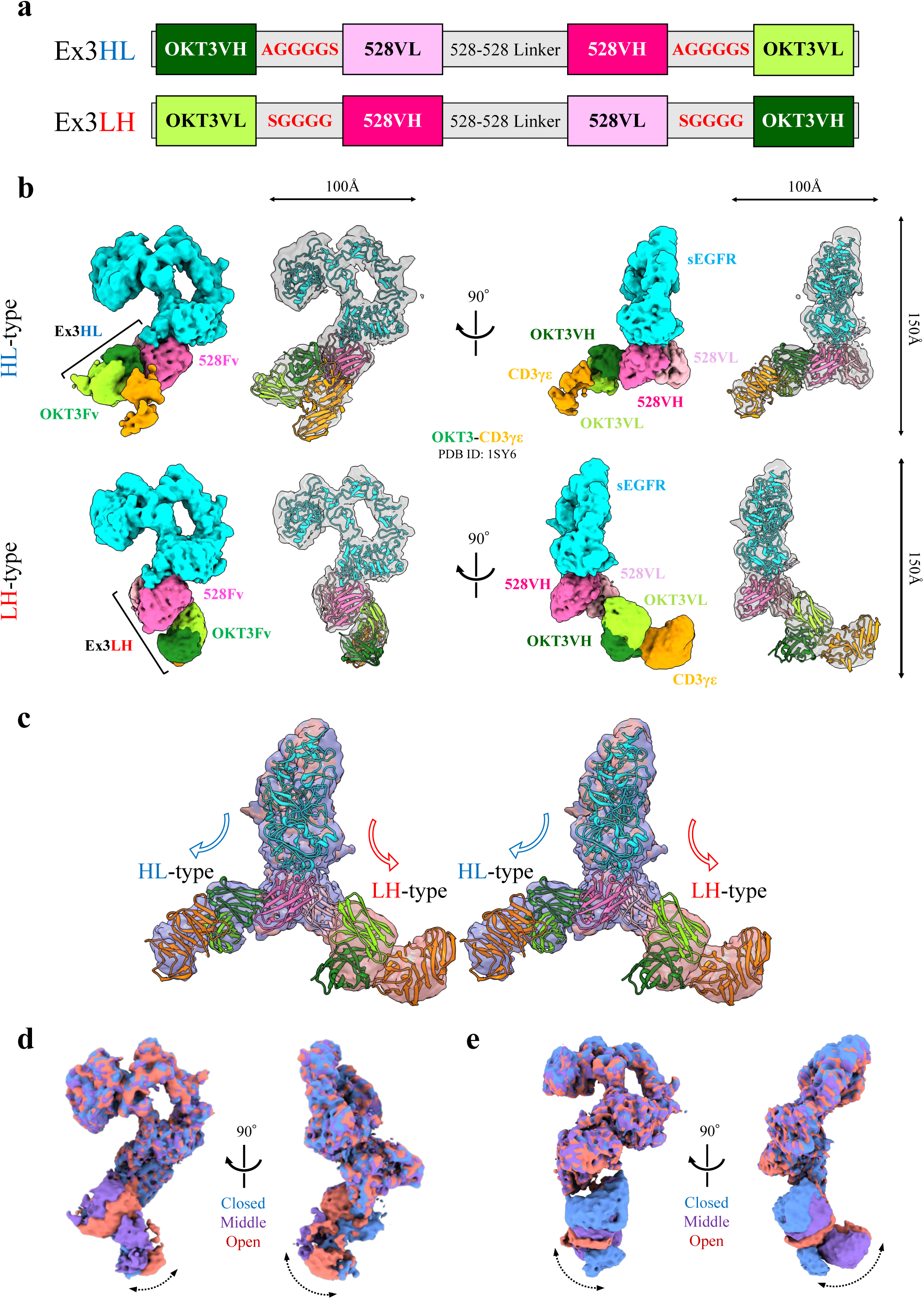
Cryo-EM structures of sEGFR-Ex3-CD3γε. (a) Domain organization of Ex3HL (upper sequence) and Ex3LH (lower). The linker regions connecting OKT3VH/VL and 528VL/VH are shown in red. (b) Overview (‘middle’ conformations, from two different angles) of the EM map (left cartoons) and rigid-body fitting (right cartoons) of sEGFR-Ex3HL-CD3γε (upper structures) and sEGFR-Ex3LH-CD3γε (lower). The structure of sEGFR-528Fv(Ex3LH) obtained by Local Refinement and the previously reported crystal structure of OKT3-CD3γε (PDB ID: 1SY6) were each fitted to the EM maps as rigid bodies. (c) Stereo view of the comparison of the HL- and LH-type rigid-body fitted structures in Fig. b. The sEGFR-528Fv domains of both types are superimposed. The HL-type map is indicated with a blue arrow and the LH-type map with a red arrow. (d, e) Superimposed EM maps of closed (blue), middle (purple) and open (red) conformations of sEGFR-Ex3-CD3γε (d, HL-type; e, LH-type). The curved dashed double arrows indicate the assumed movement of OKT3-CD3γε.

For the development of more functional drugs, the development of BsAbs is expected to continue to advance in the future and occupy an extremely important position in the pharmaceutical industry. In addition, it is desired to develop more effective drugs to treat cancer. Therefore, it is important to elucidate the mechanism of activity enhancement and apply that knowledge to establishing techniques for producing highly effective antibody drugs; and, for that, structural information is essential. However, there are few reports on structural information of BsAbs in complex with antigens, and there have been no structural comparisons of BsAbs with the same parent antibody and different domain arrangements.

In this study, we performed cryo-electron microscopy (cryo-EM) single particle analysis of both HL- and LH-type ternary complexes of Ex3 with CD3 gamma-epsilon heterodimer (CD3γε) and the soluble extracellular domain of EGFR (sEGFR). Based on the cryo-EM structures obtained, we discuss the mechanism of activity enhancement of BsAbs by domain rearrangement and also report structural information on the binding mode of 528 to EGFR, a potent anti-cancer antibody for pharmaceutical applications, in comparison with cetuximab, a major therapeutic anti-cancer antibody.

## Materials and Methods

### • Protein expression and purification

The HL- and LH-type Ex3 used in this study are humanized^15^ single-chain diabodies^16^ (hEx3-scDb). H528 VH carries a high-affinity HY52W mutation^17,18^ (Supplementary Fig. 1). For simplicity, these are referred to as just Ex3HL and Ex3LH. Also, their Fvs are referred to as OKT3Fv and 528Fv and their VHs and VLs are referred to as OKT3VH, OKT3VL and 528VH, 528VL. Ex3HL/LH was expressed using the *Brevibacillus choshinensis* HPD31-SP3 expression system as described previously^16^. The medium supernatant containing Ex3HL/LH was centrifuged at 4,500 x g for 20 min and the resulting supernatant was collected. Ammonium sulfate (60% w/v) was added at 4°C with gentle mixing and stirred for 1 hour. It was then centrifuged at 4,500 x g for 20 min and the precipitates were suspended in a small volume of phosphate buffered saline (PBS) buffer (137 mM NaCl, 2.68 mM KCl, 10 mM Na_2_HPO_4_, 1.76 mM KH_2_PO_4_). The suspension was dialyzed three times with PBS buffer and centrifuged at 15,000 x g. The supernatant was subjected to Ni-nitrilotriacetic acid (Ni-NTA) affinity chromatography and eluted with PBS buffer containing 10, 50, 150, 200, 300, and 500 mM imidazole to collect fractions containing Ex3HL/LH. The crude purified sample was subjected to size exclusion chromatography (SEC) using Superdex200 10/300 in PBS buffer to obtain purified Ex3HL/LH. sEGFR was expressed using the sEGFR high-expressing Chinese hamster ovary (CHO) cell expression system as described previously^19^. The medium supernatant containing sEGFR was subjected to Ni-NTA affinity chromatography and eluted with PBS buffer containing 10 mM, 500 mM, and 1 M imidazole to obtain purified sEGFR. CD3γε was expressed and purified using the *E. coli* BL21(DE3) strain expression system as described previously^20,21^.

Genes for EGFR domain III ND2 variants (WT, S464L, G465R, K467T, and S492R) were obtained from a synthetic gene service (GenScript). The genes were cloned into the N-terminal side of a mouse Fc gene (EGFRD3-mFc) on the pCAGEN vector for mammalian expression. An Expi293F expression system was used to produce recombinant proteins. The culture supernatants were applied onto a Protein A column (KanCapA, Kaneka) and the elution fractions were dialyzed in PBS buffer. The antigen-binding fragment (Fab) of cetuximab and 528 were prepared by papain digestion of the parental antibodies cetuximab (purchased from Merck BioPharma) and 528, respectively. Digested samples were applied onto the Protein A column, and flowthrough fractions were collected.

### • Preparation of sEGFR-Ex3-CD3γε ternary complex

Purified sEGFR and Ex3HL/LH were mixed at an equimolar molecular ratio and incubated at 4°C for 30 minutes. To purify binary sEGFR-Ex3HL/LH complexes, mixtures were subjected to SEC using Superdex200 10/300 in PBS buffer (Supplementary Fig. 2a-d). Subsequently, for CD3γε bindings for binary complexes, a 4-fold molar ratio of CD3γε was added to the sEGFR-Ex3HL/LH complex obtained from SEC. These solutions were further incubated at 4°C for 30 min. After incubation, it was immediately used for grid preparation for cryo-EM. The final concentration of sEGFR-Ex3HL/LH was adjusted to 1∼3 μM.

### • Cryo-EM sample vitrification and data collection

Ternary complex sEGFR-Ex3HL/LH-CD3γε (3 µL) was applied to Quantifoil R1.2/1.3 Cu girds (Quantifoil) that had been glow discharged at 10 mA for 40 seconds with a sputter coater (JEOL JEC-3000FC). Samples were then vitrified using a Vitrobot Mark IV (Thermo Fisher) at 4°C and 100% humidity. Cryo-EM data were acquired using a CRYO ARM 300 II (JEOL) operated at 300 kV acceleration voltage. Movie micrographs were collected at 60,000x nominal magnification (corresponding to a physical pixel size of 0.788 Å) with a total dose of 60 electrons per Å^2^ using a K3 Camera direct electron detector (Gatan) with the SerialEM program^22^. Grid vitrification and data collection were performed under three different conditions for each of the HL- and LH-types and were used for subsequent data processing (Supplementary Table 1).

### • Data processing

A total of 9,800 movies for sEGFR-Ex3HL-CD3γε and 19,350 movies for sEGFR-Ex3LH-CD3γε were obtained by merging the three different data sets for the three different conditions (Supplementary Table 1). All data processing for each merged dataset was performed using CryoSPARC^23^ version 4.4 (Supplementary Fig. 3a-f, 4a-g). Briefly, Patch Motion Correction and Patch CTF Estimation were performed on each dataset, and then Manually Curate Exposures was used to eliminate bad micrographs by thresholding and manual selection. Particle picking was performed on the selected micrographs using Template Picker, extracted with a box size of 384×384 pixels, and then 2D Classification was performed iteratively to select particles. Ab-Initio Reconstruction and Heterogeneous Refinement were performed on the purified particles in two classes. The class with clear sEGFR characteristics was selected for Non-uniform Refinement^24^ and the initial map was obtained. Focused 3D Classification was performed for the low-density OKT3Fv-CD3γε region and three conformational (closed, middle, open) ternary complex maps were obtained, differentiated by the degree of opening of the hinge region consisting of two linkers connecting the two Fvs (here referred to as the diabody hinge (Db-hinge) region). After Reference Based Motion Correction was performed on the particles in each conformation, Non-uniform Refinement was performed to obtain three final maps. The particles of these three maps were used for 3D Variability Analysis^25^ (3DVA) to observe the flexibility of the OKT3Fv-CD3γε region (Supplementary Video 1, 2). In addition, Local Refinement was performed on the sEGFR-528Fv region of the initial map of the LH-type after Reference Based Motion Correction to obtain a final local map for modeling (sEGFR-528Fv(Ex3LH)). The signal was subtracted from the OKT3-CD3γε region by Particle Subtraction. All the final maps obtained were sharpened using Sharpening Tools and the local resolution was estimated using Local Resolution Estimation.

### • Model building

The crystal structure of sEGFR-GC1118A^26^ (PDB ID: 4UV7) was used for sEGFR of the initial model of the sEGFR-528Fv(Ex3LH) complex. The Ex3LH structure predicted by AlphaFold2^27^ was used for 528Fv of the initial model. These structures were rigid-body fitted to the local map of sEGFR-528Fv(Ex3LH) using UCSF ChimeraX^28^. Missing loops and glycans in low-density regions were manually deleted, and several rounds of manual adjustment in COOT^29^ and refinement using Phenix.real_space_refine^30^ were performed to obtain the final structure (Supplementary Table 2). This sEGFR-528Fv(Ex3LH) model and the known crystal structure of OKT3-CD3γε^21^ (PDB ID: 1SY6) were each rigid-body fitted to the ternary complex maps. The structure figures and movies in this article were generated using PyMOL (Schrödinger, LLC.), UCSF Chimera^31^ and UCSF ChimeraX.

### • Biolayer interference (BLI) measurements

Interactions between the EGFRD3ND2-mFc variants and Fabs were evaluated by the biolayer interference (BLI) measurements using Octet N1 (Sartorius). Anti-mouse Fc capture (AMC) biosensors were used to capture the EGFRD3ND2-mFc variants, whose concentrations were set to 0.4μM ∼ 1μM (Supplementary Table 3, 4). Experiments were performed in PBS buffer at room temperature. The concentrations were set to 200 nM for cetuximab Fab and 125 nM for 528 Fab.

## Results and Discussion

### • Cryo-EM structures of sEGFR-Ex3-CD3γε

The cryo-EM structures of the sEGFR-Ex3-CD3γε ternary complex were determined at resolutions of 3.64 Å for the HL-type and 3.29 Å for the LH-type. In both structures, densities corresponding to sEGFR and 528Fv were clearly observed, whereas the OKT3Fv and CD3γε regions represented relatively low resolution due to their inherent flexibility. An atomic model was built for the sEGFR-528Fv region of the LH-type using the local map from Local Refinement on the sEGFR-528Fv region to understand the molecular details. To understand the global topology of the ternary complexes, the atomic models of sEGFR-528Fv(Ex3LH) and previously reported crystal structure of OKT3-CD3γε^21^ (PDB ID: 1SY6) were rigid-body fitted into the cryo-EM maps (Fig. 1b). Both sEGFR-Ex3HL-CD3γε and sEGFR-Ex3LH-CD3γε were found to have dimensions of approximately 100 Å x 100 Å x 150 Å and an arcuate shape. 528Fv (pink) was bound to domain Ⅲ of sEGFR (cyan) (details of the binding mode are described below); OKT3Fv-CD3γε (green and orange) was located on the opposite side of the antigen-binding region from 528Fv. Although the density of the linkers connecting 528Fv and OKT3Fv was obscure, the N- and C-terminus of each Fv expected to be connected by the linker were located proximally, demonstrating the validity of the structure. Comparing the two structures, sEGFR-528Fv and OKT3Fv-CD3γε had similar shapes in both structures, but the relative positions of sEGFR-528Fv and OKT3Fv-CD3γε were different. Consequently, the overall structure had an inverse curve around the Db-hinge region consisting of two linkers connecting 528Fv and OKT3Fv. Superposition of sEGFR-528Fv revealed that the position of OKT3 relative to 528Fv is opposite between HL- and LH-types, rendering the relative position between CD3γε and sEGFR quite different (Fig. 1c). This difference in the relative position of two antigens can explain the 1000-fold larger change in anticancer activity (discussed further below). It is noted that this marked change in the relative position of the two antigens occurred just by changing the order of the Fv domains. In both HL- and LH-types, the antigen-binding regions of the two Fvs face away from each other and consequently there is no steric hindrance between sEGFR and CD3γε. This is consistent with the previous report showing that both Ex3HL and Ex3LH can bind to their soluble antigens with affinities similar to their respective parent antibodies, and, moreover, they can bind to one soluble antigen without decreased affinity even when complexed with the other^32^.

A series from 3D Classification yielded cryo-EM maps of the three conformations of the sEGFR-Ex3-CD3γε ternary complex for each of the HL- and LH-types (Supplementary Figs. 3c, 4c). These three conformations are distinguished by the degree of opening of the Db-hinge region and are referred to as ‘closed,’ ‘middle,’ and ‘open’. The global resolution of the HL-type final map was 3.91 Å for the closed, 3.64 Å for the middle, and 3.85 Å for the open conformation. Those of the LH-type were 3.40 Å for closed, 3.29 Å for middle, and 3.31 Å for open. Resolution in the OKT3Fv-CD3γε region is relatively low, 4∼6 Å (Supplementary Fig. 3d-f, 4e-g), and superposition of the three conformations revealed fluctuations in this region for both HL- and LH-types (Fig. 1d, 1e). To further observe these fluctuations in detail, we performed 3DVA^25^ and observed continuous motion in both HL- and LH-types, showing flexibility starting from the Db-hinge region (Supplementary Video 1, 2).

To conclude, differences between the HL- and LH-types were revealed, specifically: (i) the relative positions of the two Fvs change drastically depending on the order of the Fv domains (Fig. 1c); (ii) the relative positions of the antigens consequently change significantly; and (iii) Db-hinge motion between the two Fvs allows for flexible movement. Understanding these large differences in the relative positioning of Fvs and/or antigens, and the flexibility between Fvs, provides a sound basis to explain the differences in anti-cancer activity between the HL- and LH-types.

### • Structural basis for Ex3 activity enhancement by domain rearrangement

The difference between the HL- and LH-types is the order of the connected domains, which in turn is a property of changing the position of the linker that connects the two domains. The Db-hinge region is formed by the two OKT3VH/VL – 528VL/VH linkers (red in Figs. 1a, 2a, 2b) and is the most important determinant of the overall structure. In the HL-type, the C-terminus of OKT3VH is linked to the N-terminus of 528VL, and the C-terminus of 528VH is linked to the N-terminus of OKT3VL. In the LH-type, the C-terminus of OKT3VL is connected to the N-terminus of 528VH, and the C-terminus of 528VL is connected to the N-terminus of OKT3VH. The structures of Ex3 show that the two linkers of OKT3VH/VL – 528VL/VH are located in opposite positions in the HL- and LH-types (Figs. 2a, 2b), which explains why their direction of bending was different. As observed in the 3DVA^25^, the two Fvs are expected to be connected with substantial open-closed flexibility (Supplementary Videos 1, 2).

**Figure 2:**
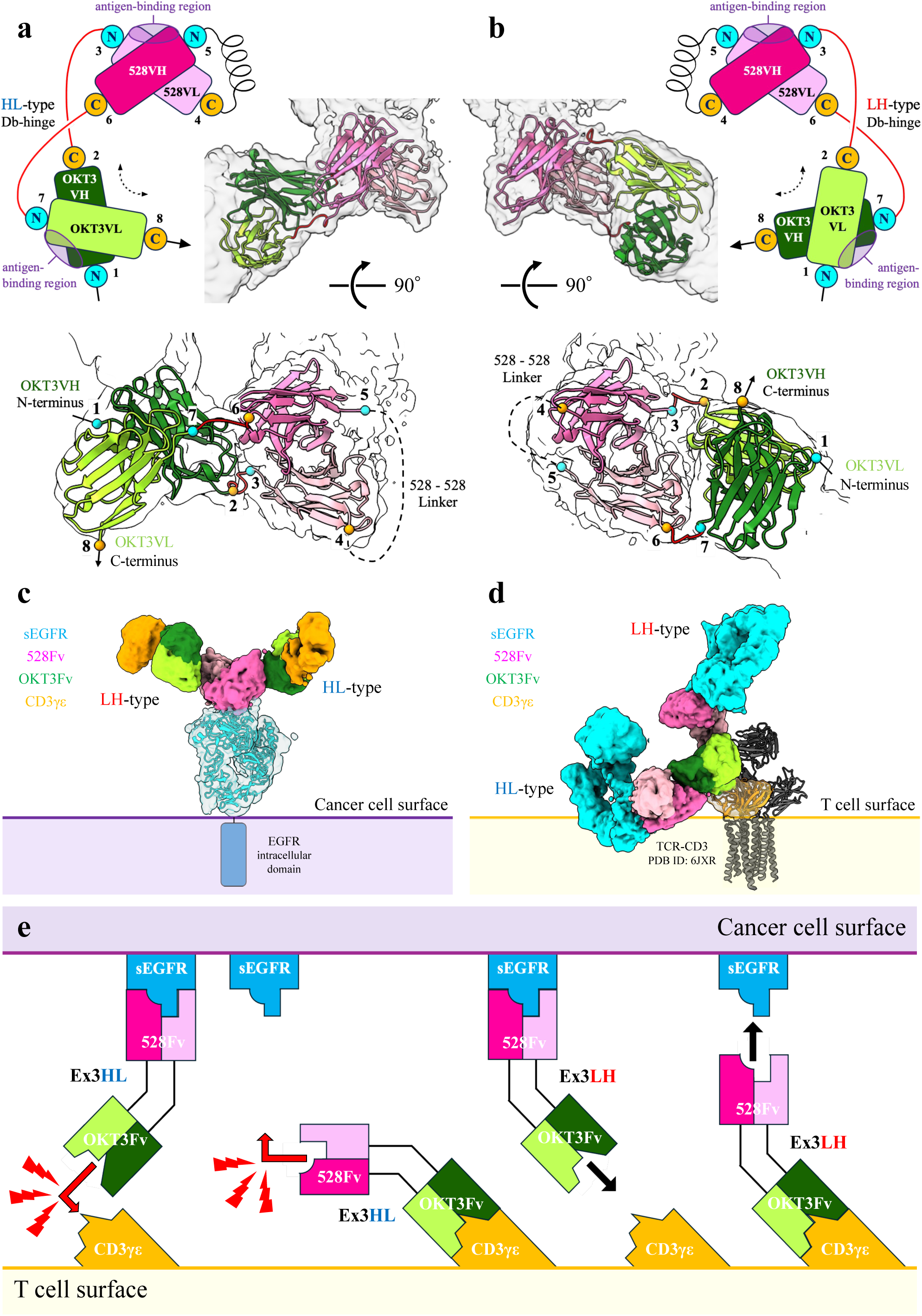
Structural basis for Ex3 activity enhancement by domain rearrangement. (a, b) Schematic diagram and space models (enclosing secondary structure) of Ex3 for (a) the HL-type and (b) the LH-type. The positional relationship between 528Fv and OKT3Fv of Ex3HL/LH predicted by AlphaFold2 was corrected and fitted to the middle conformation map. The N-terminus of each domain is shown as a cyan circle, and the C-terminus as an orange circle. Numbering 1∼8 indicates the order of connection in the sequence. The space models are illustrated from two viewing angles. (c) Predicted binding mode of Ex3HL/LH on the cancer cell surface. Both HL- and LH-type middle conformation maps were fitted to the sEGFR model. (d) Predicted binding mode of Ex3HL/LH on the T cell surface. Both HL- and LH-type middle conformation maps were fitted to CD3γε of the TCR-CD3 complex (PDB ID: 6JXR). (e) Schematic diagram of bridging between T cells and cancer cells by Ex3HL (left side) or Ex3LH (right side) when attached first to sEGFR on the cancer cell (left) or CD3γε on the T cell (right).

Obviously, a change in the orientation between the Fvs in Ex3 changed the position of the antigen-binding regions, as well as the mode of bridging between the two antigens. Here, we predicted the binding mode of Ex3HL/LH on the cancer cell surface based on a previous report on the dynamics of sEGFR on the cell surface^33^ (Fig. 2c). In both HL- and LH-types, although their orientation differs, OKT3-CD3γε is oriented toward the surface of the cancer cells, suggesting that in both HL- and LH-types the OKT3 Fv can bind to CD3γε even when Ex3 is bound to the cancer cell surface. Similarly, we predicted the binding mode of Ex3HL/LH on the T cell surface by superimposing CD3γε on the known cryo-EM structure of TCR-CD3^34^ (PDB ID: 6JXR) (Fig. 2d). It can be seen that, in the LH-type, 528Fv-sEGFR is oriented vertically to the T cell surface, whereas in the HL-type, it is oriented horizontally. Ex3LH oriented vertically on the T cell appears to be able to bind to sEGFR, whereas Ex3HL would cause steric hindrance between sEGFR and the lipid bilayer of the T cell (Fig. 2d). This was the finding for all conformations from closed to open. In a previous report using flow cytometry, it was demonstrated that both Ex3HL and Ex3LH bound to the cancer cell were able to bind to purified CD3γε, and Ex3LH bound to the T-cell was able to bind to purified sEGFR However, Ex3HL bound to T cells was unable to bind to purified sEGFR^11^. Our cryo-EM structures clearly explain these observations (Fig. 2d): that is, sEGFR bound to Ex3HL on the T cell collides with its lipid bilayer; whereas Ex3LH on the T cell, and both Ex3HL and Ex3LH on the cancer cell, have enough space for binding to the other antigen.

Based on the structures revealed here, the characteristics of Ex3-induced bridging between cancer cells and T cells can now be explained (Fig. 2e). When Ex3HL/LH bridges cancer cells and T cells, there are two possible patterns: Ex3HL/LH first binds to cancer cells and then captures T cells, or first binds to T cells and then captures the cancer cells. The difference between Ex3HL and Ex3LH is represented by the difference in their angles of bending. Regardless of the order of cell binding, Ex3LH is expected to be able to bridge cancer cells to T cells because the antigen-binding region of the remaining Fv is open to the other antigen. However, in both patterns of Ex3HL, because the antigen-binding region of the remaining Fv and the epitope of the antigen are positioned at a bend, and steric hindrance between the cancer cell and the T cell is expected to occur, which would prevent bridging of the cells. Therefore, because Ex3LH can avoid any steric hindrance between cells, it has a higher bridging capacity than Ex3HL, explaining its consequently higher anti-cancer activity. This both confirms and explains a previous report, using atomic force microscopy (AFM), that Ex3LH bridges between cancer cells and T cells more strongly than Ex3HL^12^. Comparing the two bridging patterns of Ex3HL, when Ex3HL binds to T cells first, the antigen-binding region of 528Fv is oriented horizontally and is not exposed to the outside of the cell, which would make contact with the other cell even more difficult compared with when Ex3HL binds to cancer cells first. In addition, the extracellular domain of EGFR is dynamic^33^, so it may be more likely to avoid steric hindrance when it first binds to cancer cells. This means that even when bridging with the same type of BsAb, there may be differences in its bridging ability depending on the cell to which it binds first. Thus, when considering bridging by BsAbs, the binding angle and flexibility of the BsAb itself, the angle of the epitopes of both antigens to the cell surface, and even their dynamics on the cell surface must be considered in an integrated manner.

Taking these observations together, the domain rearrangement of BsAbs results in a significant change in the relative angle between the two Fvs and even between the antigens. Depending on the binding angle of the two cells, some cause steric hindrance (the HL-type in the case of Ex3), while others that avoid steric hindrance (the LH-type in the case of Ex3) can exhibit effective bridging ability. The difference in the bridging ability results in the difference in activity of BsAbs. This avoidance of steric hindrance is considered to be the fundamental mechanism for enhancing BsAb activity through domain rearrangement. In addition, binding angle has a significant effect on the activity of BsAbs, and domain order was found to be extremely important for enhancing activity since it determines the relative position between the two Fvs and the two antigens.

### • Cryo-EM structure of sEGFR-528Fv(Ex3LH)

In both HL- and LH-types, the structure of sEGFR-528Fv was revealed in high resolution using cryo-EM, providing details of the binding mode of 528Fv for sEGFR. Local Refinement of the sEGFR-528Fv region with a focus mask for the initial map of sEGFR-Ex3LH-CD3γε, yielded a map with a global resolution of 2.93 Å (Supplementary Fig. 4c). Although the local resolution of domain Ⅰ and domain Ⅳ of sEGFR is relatively low due to their flexibility (Supplementary Fig. 4d), most of the regions retain the density quality allowing the assignment of amino acid residues, building the local structure of sEGFR-528Fv(Ex3LH) (Fig. 3a). This structure confirms the previous report^35^ of the epitope of EGFR to which 528 binds and provides further details of its binding mode. 528 captured loop 353-359 of sEGFR, several of the amino acids of which were found to form hydrogen bonds (Fig. 3b). The following amino acids were found to be important in sEGFR-528 binding: R353, D355, S356, and H359 of sEGFR; S31, W33, S95, G96, and Y99 of 528VH; and G91 of 528VL. Furthermore, W52 and W33 of 528VH are close to L325 of sEGFR, suggesting that van der Waals interactions may contribute to the binding (Fig. 3c). This result also explains the increased affinity of 528 due to the Y52W mutation on VH^17,18^.

**Figure 3:**
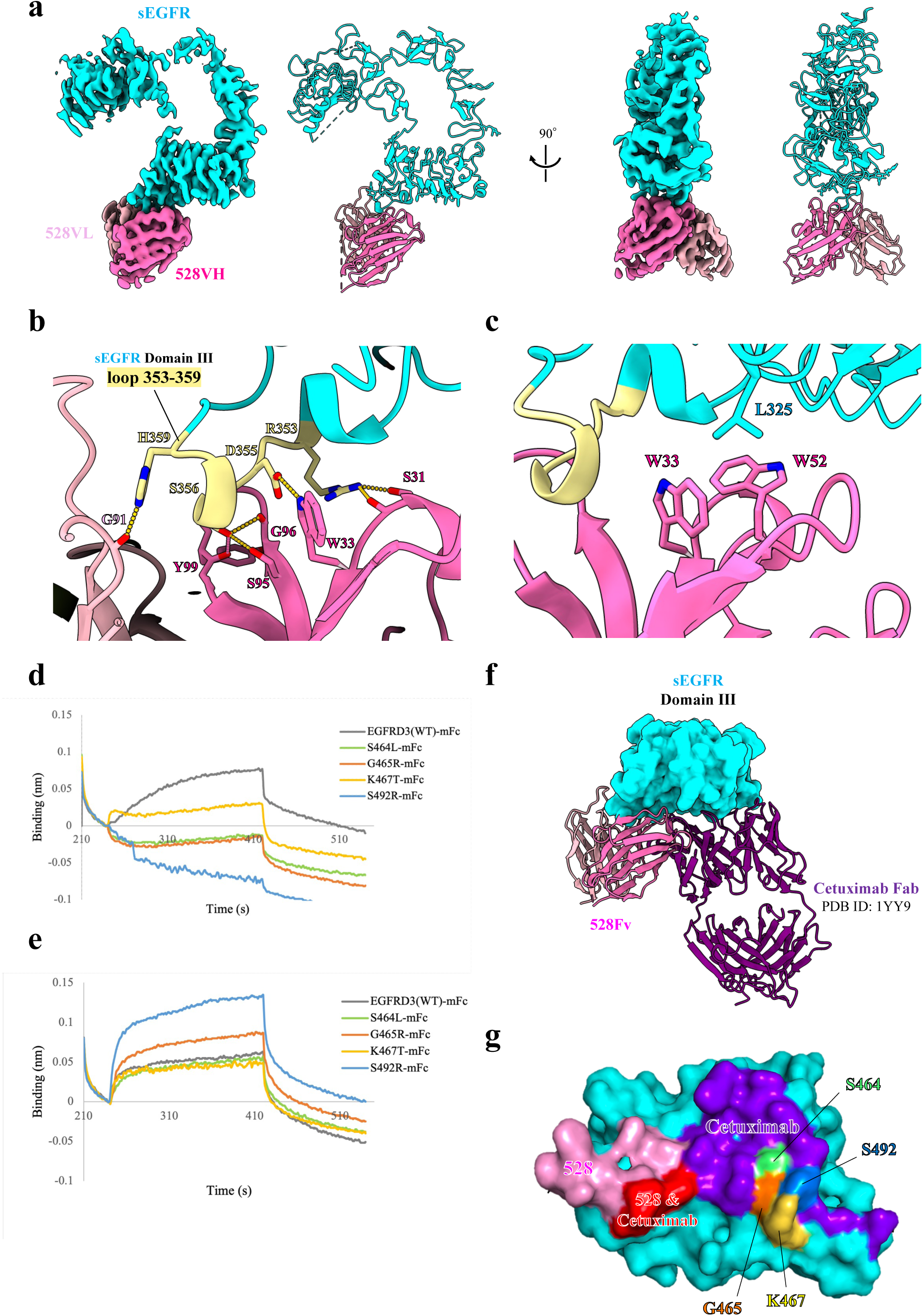
Cryo-EM structure of sEGFR-528Fv(Ex3LH). (a) Overview of the EM map from Local Refinement and coordinates of sEGFR-528Fv(Ex3LH). sEGFR is shown in cyan, 528VH in pink, and 528VL in light pink. (b) Close-up view of the interface between sEGFR and 528Fv. The hydrogen bonds are indicated by straight segmented yellow bars. (c) Relative position of W52 and W33 of 528VH and L325 of sEGFR. (d) Interactions between cetuximab Fab and EGFRD3-mFc for each mutation, analyzed by BLI. (e) Interactions between 528 Fab and EGFRD3-mFc for each mutation, analyzed by BLI. (f) Comparison of the binding modes of 528 and cetuximab to domain Ⅲ of sEGFR, shown as a surface model. (g) Surface model showing binding sites on the sEGFR domain Ⅲ surface: pink, 528; purple, cetuximab; and red for the overlapping region between the two. The position of each mutation is shown in green for S464L, orange for G465R, yellow for K467T, and blue for S492R.

EGFR is a major cancer antigen, and therapeutic anti-cancer antibodies have been developed using anti-EGFR antibodies such as cetuximab. An important issue with therapeutic anti-EGFR antibodies is their reduced efficacy due to EGFR mutations. Therefore, it is important to use EGFR antibodies that are tailored to each patient’s EGFR mutation. Furthermore, it is important to know the epitope differences between each antibody in detail. In the present study, based on knowledge gained about the high-resolution structure of sEGFR-528Fv, we discuss the pharmacological significance of 528 to cetuximab, which is in clinical use. We evaluated the interaction of 528 and cetuximab with four cetuximab-resistant sEGFR domain Ⅲ mutants (S464L, G465R, K467T, and S492R) ^36,37^ using BLI measurements. Cetuximab lost binding ability in the presence of all these mutations (Fig. 3d), whereas 528 retained binding ability to all of them (Fig. 3e). The epitope region of 528 in domain Ⅲ of sEGFR is shifted with respect to that of cetuximab^38^ (PDB ID: 1YY9) (Fig. 3f). All residues substituted in the present study are located around the epitope of cetuximab, but not in the epitope of 528 (Fig. 3g), which reasonably explains the results of the BLI interaction analysis (Fig. 3d, 3e). However, cetuximab and 528 were found to share parts of their epitopes (Fig. 3f, 3g), which explains the previously reported competitive binding of cetuximab and 528 to sEGFR^35^. These results indicate that 528 binds to domain Ⅲ of sEGFR in a similar way to cetuximab while 528 can avoid the reported cetuximab-resistant mutations, which suggests that 528 could be used as a second line of attack replacing cetuximab treatment where these mutations are present.

## Conclusion

This study clarified the structural differences between HL- and LH-type Ex3 BsAb using cryo-EM single-particle analysis and obtained structural insights into the activity enhancement mechanism resulting from domain rearrangement. It was found that the order of linkage of the Fv domains causes a significant difference in the relative position of the antigens, leading to the ability of cells to make contact with each other, or their inability to do so due to steric hindrance. Furthermore, the pharmacological significance of 528 to cetuximab, which is in clinical use, was also clarified from a structural point of view. This study bridges the previously existing gap between molecular studies and cell biological findings.

Since the basic structure of antibodies is highly conserved, the results of this study are expected to be applicable to BsAbs other than Ex3. Evaluating the effect of domain rearrangement is likely to become increasingly important as the development of novel therapeutic antibodies through protein engineering flourishes, including BsAbs in many different formats^18,39^ and multispecific antibodies^40,41^ (MsAbs), which are larger in size and more complicated. Furthermore, in addition to the HL- and LH-types discussed here, there are many other types of domain rearrangements of BsAb^16^ such as tandem single-chain Fvs (taFvs). If the relationships between domain rearrangements, linker configurations, and structures of all types can be generalized, and the optimal construct for each can be predicted with high accuracy, the process of developing new highly functional BsAb- and MsAb-based drugs will be greatly accelerated. In this sense, the methods and findings reported here provide a stepping stone towards establishing a design foundation for BsAb (and even MsAb) development.

---

**Supplementary Figure 1:**
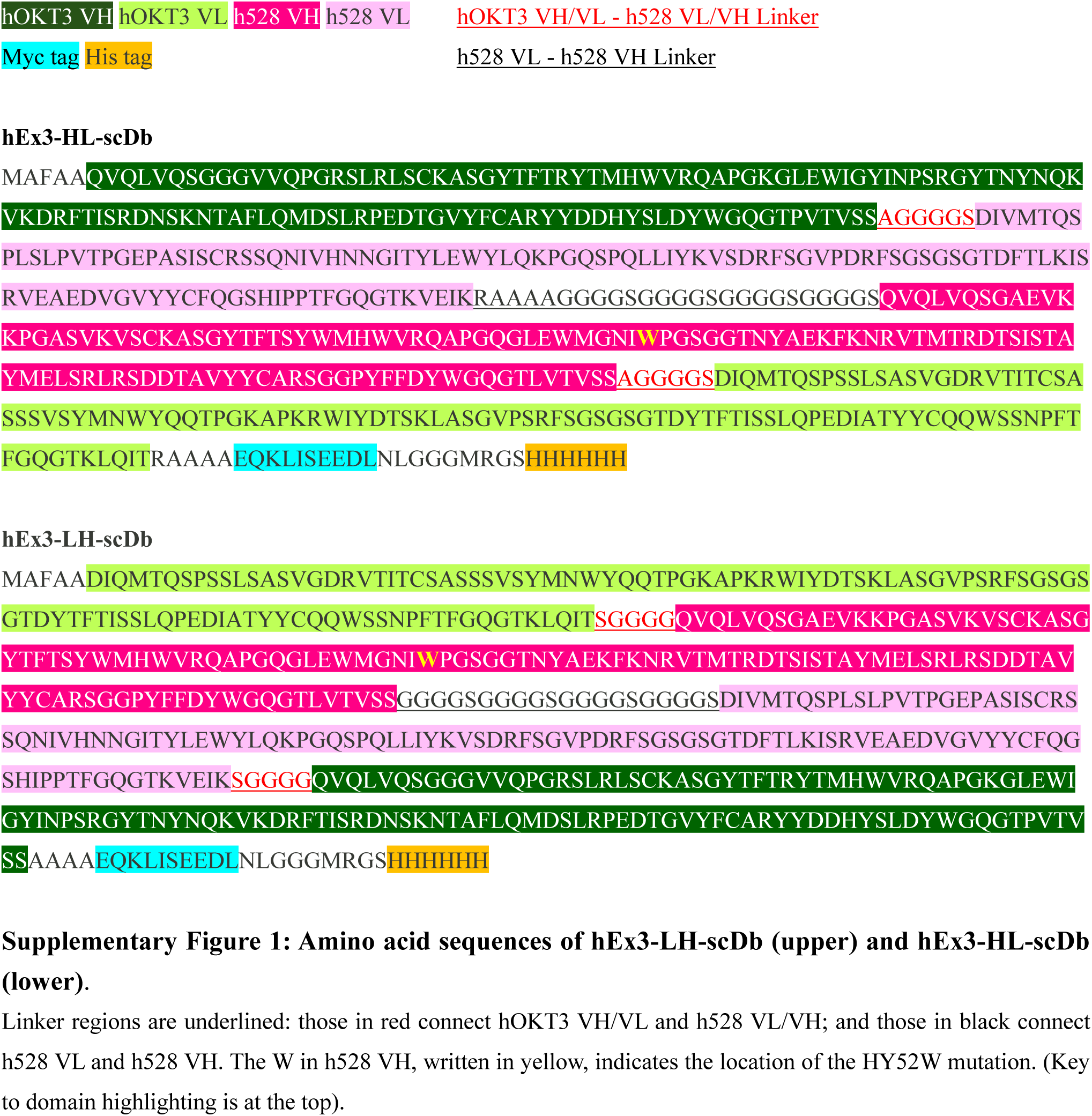
Amino acid sequences of hEx3-LH-scDb (upper) and hEx3-HL-scDb (lower). Linker regions are underlined: those in red connect hOKT3 VH/VL and h528 VL/VH; and those in black connect h528 VL and h528 VH. The W in h528 VH, written in yellow, indicates the location of the HY52W mutation. (Key to domain highlighting is at the top).

**Supplementary Figure 2:**
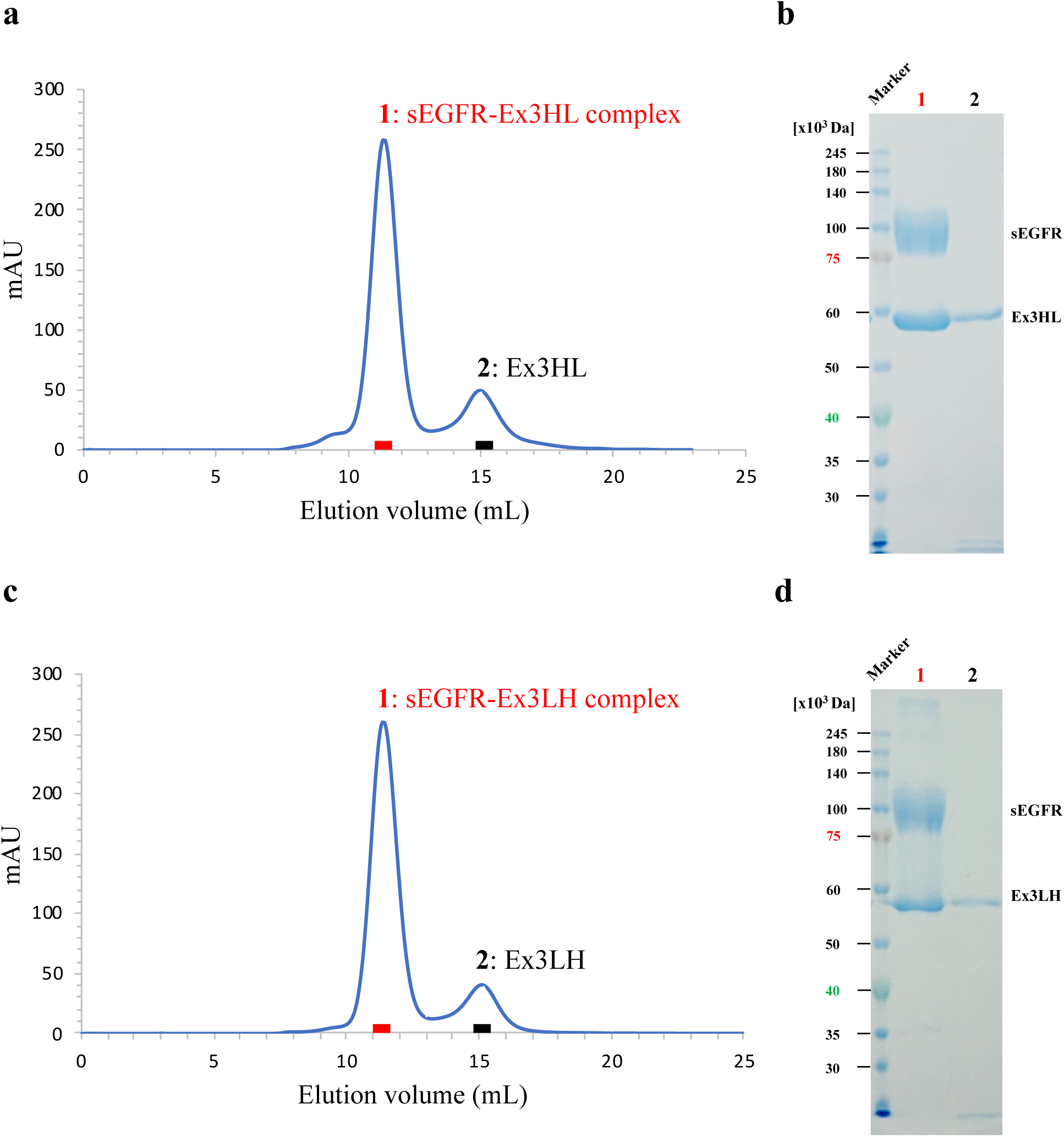
Purification of sEGFR-Ex3 complex by Superdex 200 10/300 GL size-exclusion chromatography (SEC). (a) SEC chromatogram for sEGFR-Ex3HL. (b) SDS-PAGE of each SEC peak for sEGFR-Ex3HL. (c) SEC chromatogram for sEGFR-Ex3LH. (d) SDS-PAGE of each SEC peak for sEGFR-Ex3LH.

**Supplementary Figure 3:**
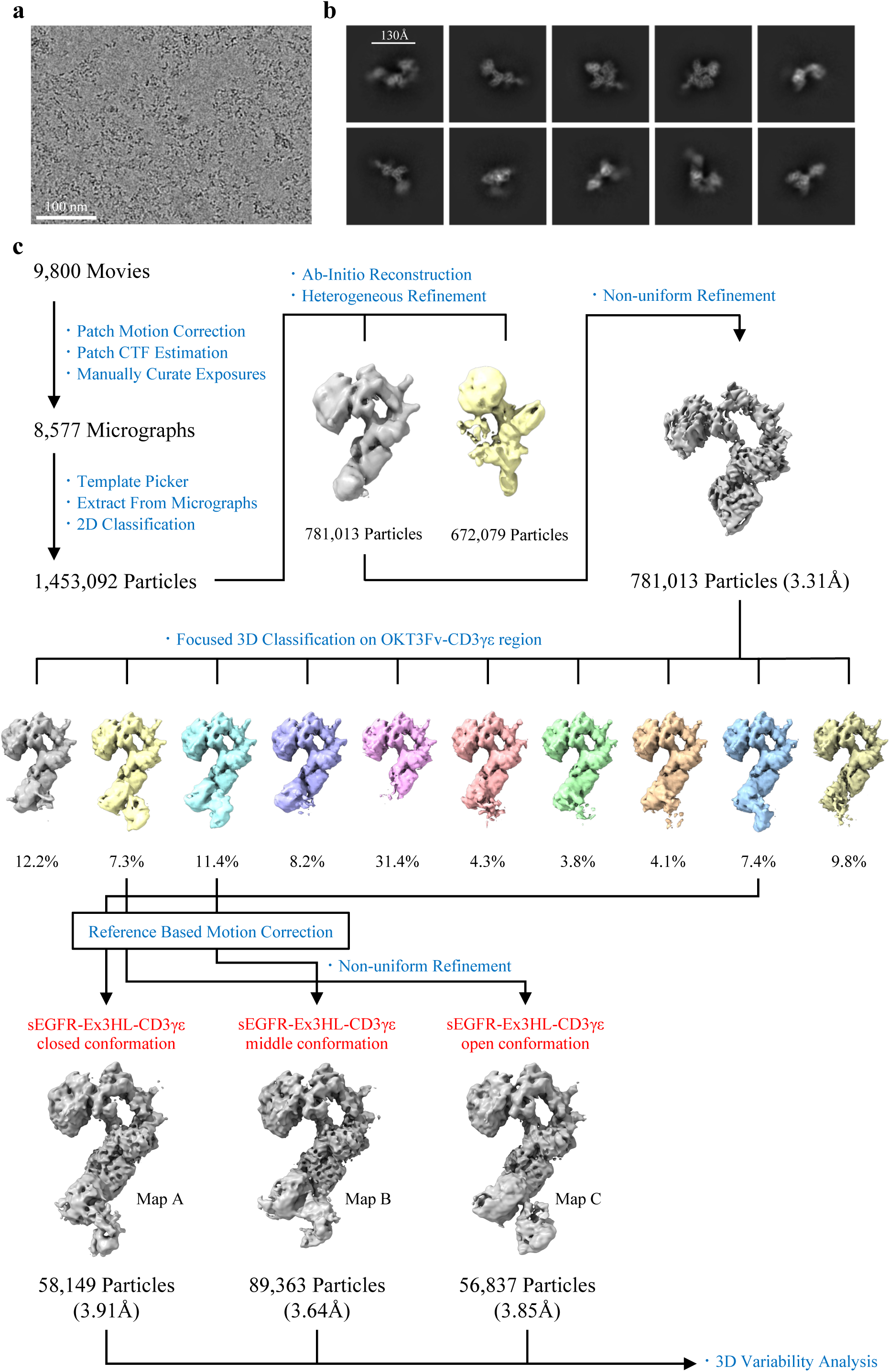
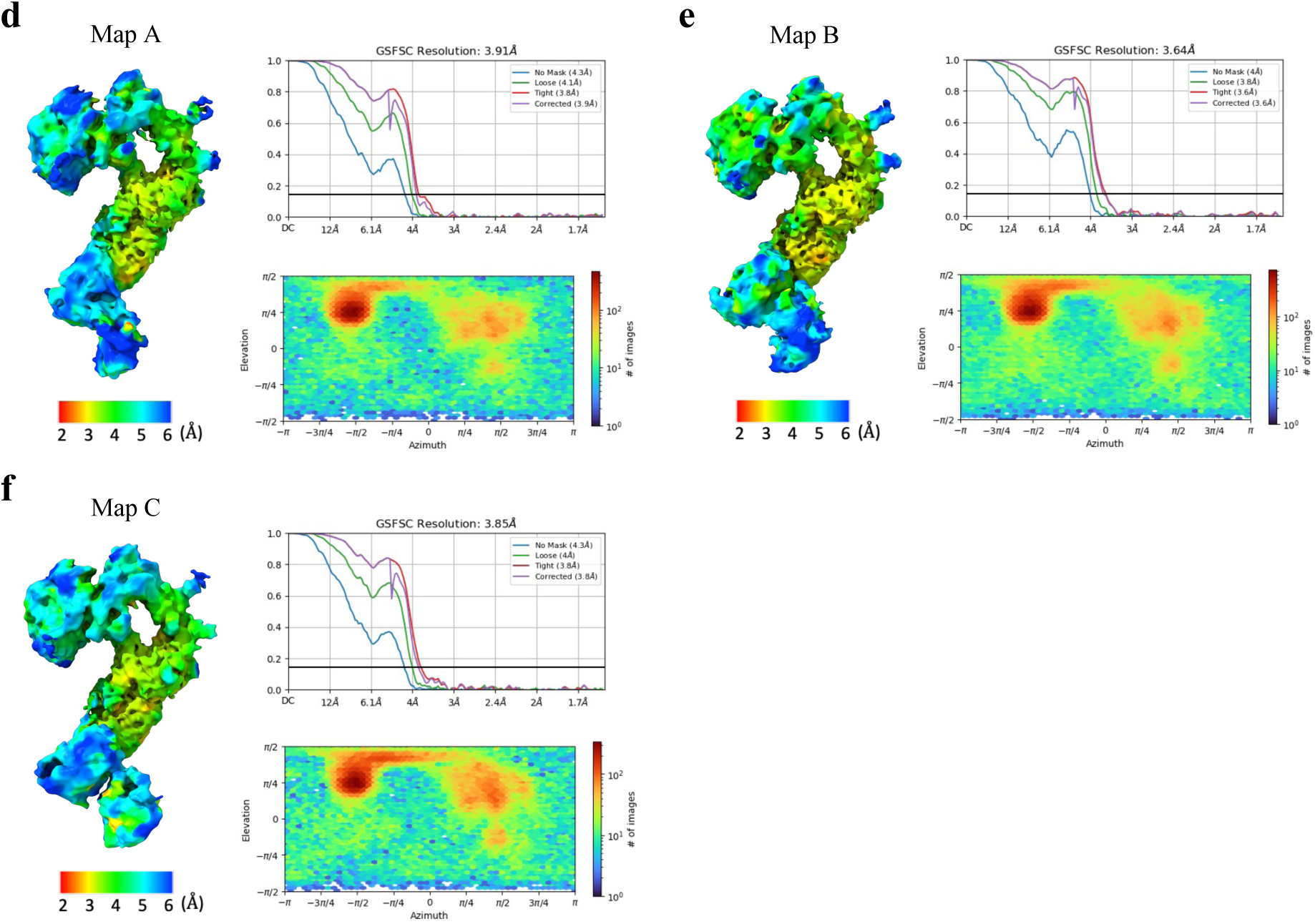
Cryo-EM data processing of sEGFR-Ex3HL-CD3γε structure. (a) A representative cryo-EM micrograph of sEGFR-Ex3HL-CD3γε. (b) Representative 2D class averages of sEGFR-Ex3HL-CD3γε. (c) Cryo-EM data processing workflow. Focused 3D Classification was performed on the low-density OKT3Fv-CD3γε region and three maps were refined, differentiated by the degree of opening of the Db-hinge region (maps A, B, C). (d, e, f) FSC, local resolution, and angular distribution for the final maps A, B, and C, respectively.

**Supplementary Figure 4:**
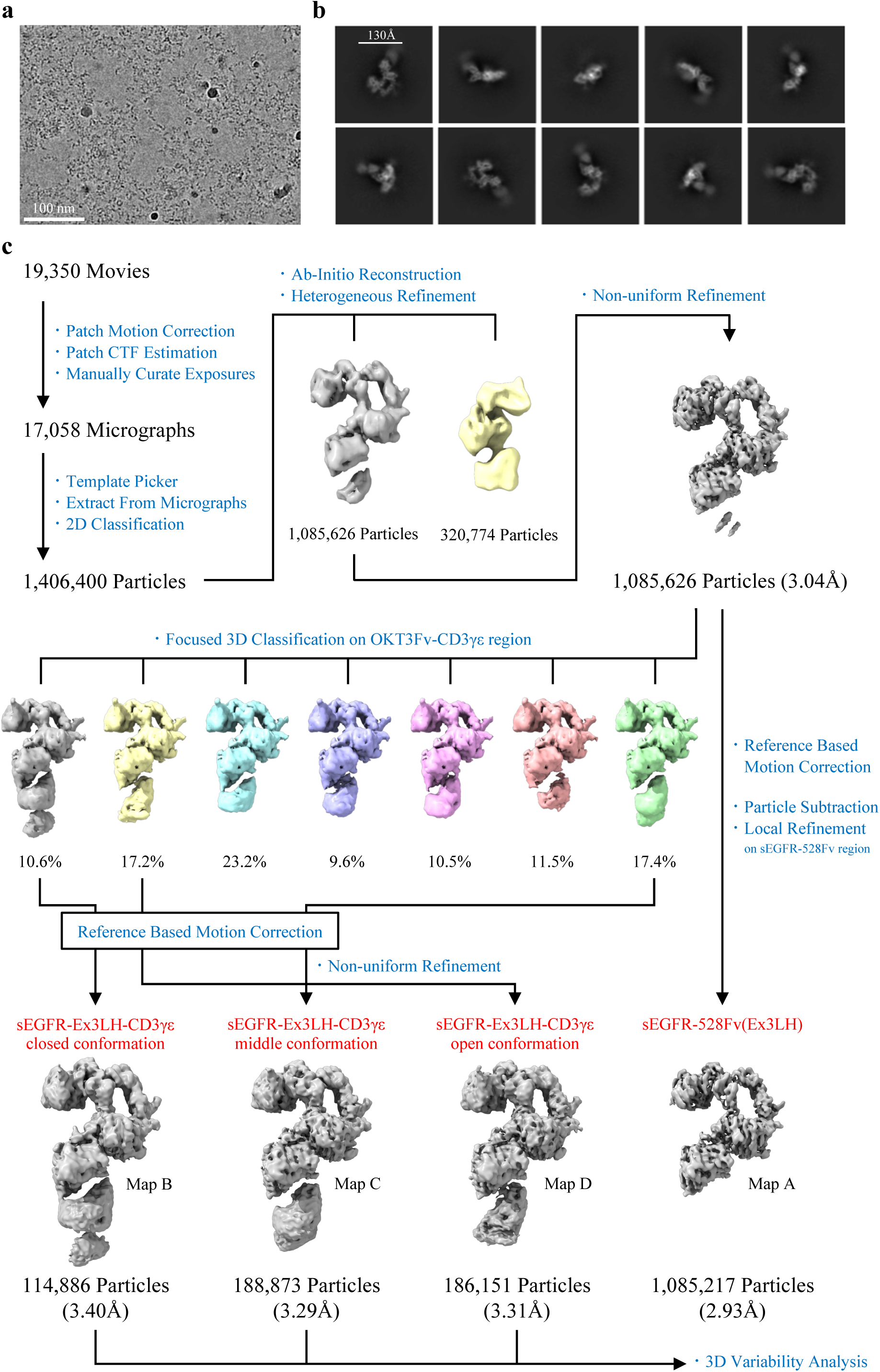
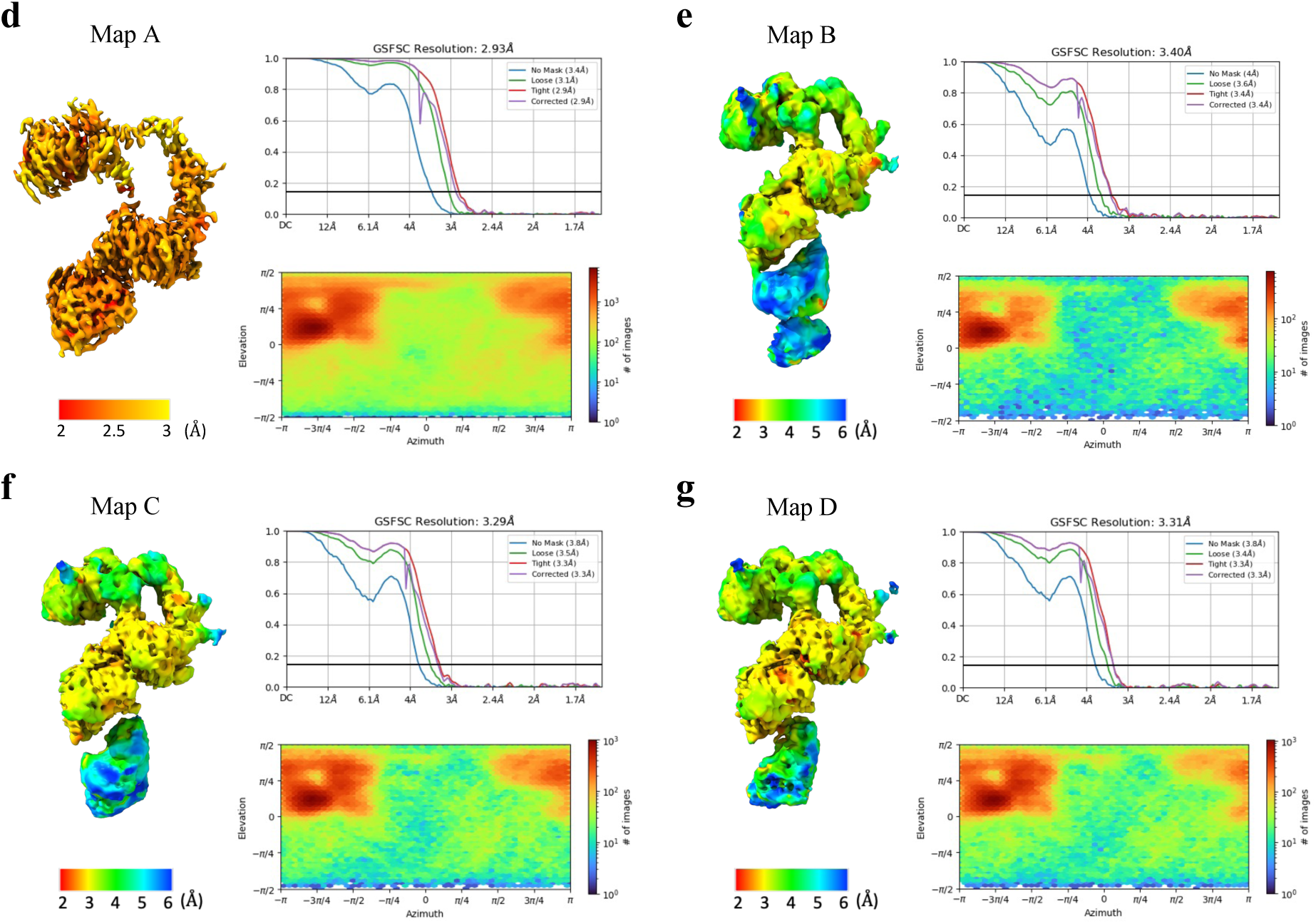
Cryo-EM data processing of sEGFR-Ex3LH-CD3γε structure. (a) A representative cryo-EM micrograph of sEGFR-Ex3LH-CD3γε. (b) Representative 2D class averages of sEGFR-Ex3LH-CD3γε. (c) Cryo-EM data processing workflow. Local Refinement was performed on the sEGFR-528Fv(Ex3LH) region and a local map was obtained for modeling (map A). Focused 3D Classification was performed on the low-density OKT3Fv-CD3γε region and three maps were refined, differentiated by the degree of opening of the Db-hinge region (maps B, C, D). (d, e, f, g) FSC, local resolution, and angular distribution for the final maps A, B, C, and D, respectively.

**Supplementary Table 1:**
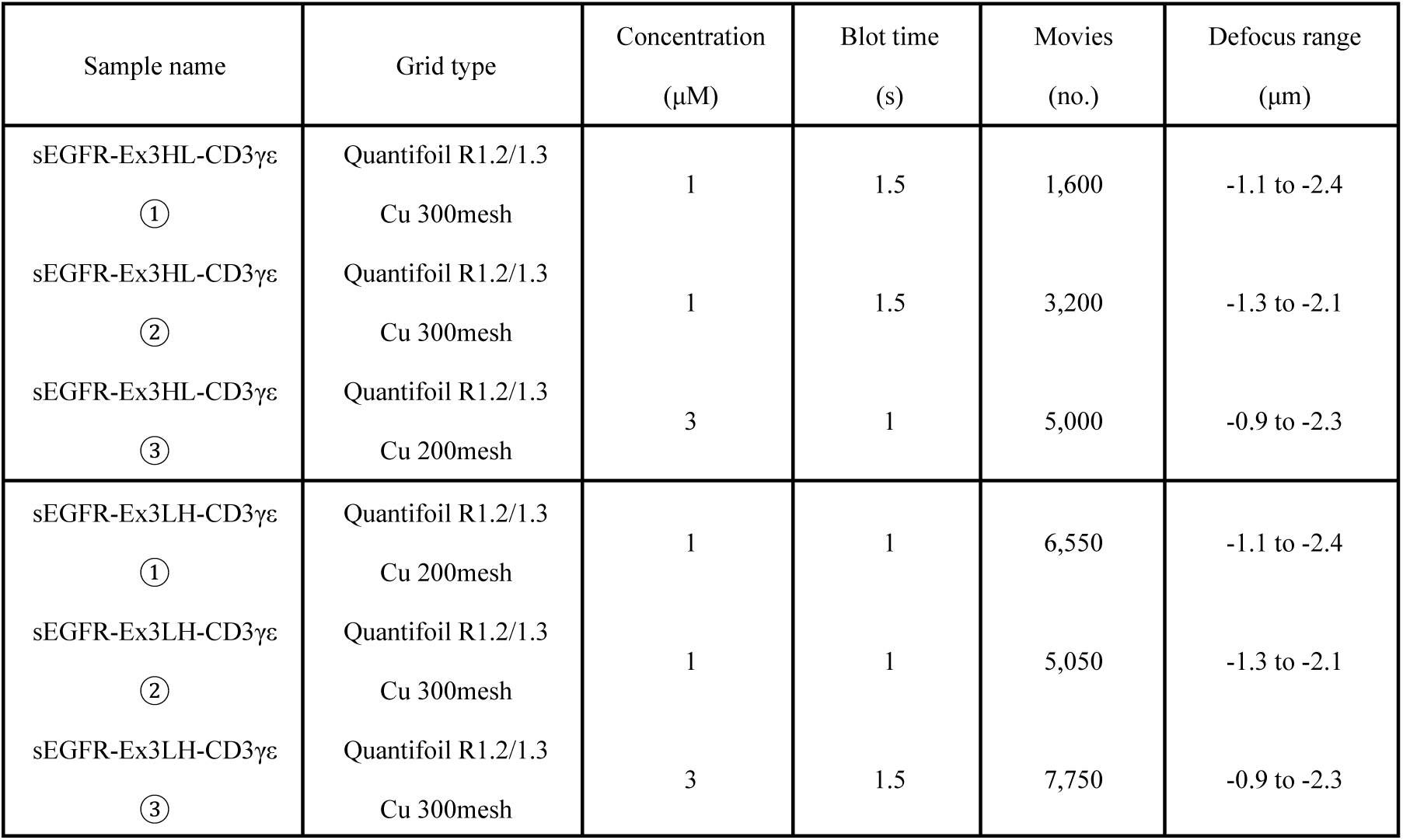
Details of cryo-EM data collection.

**Supplementary Table 2:**
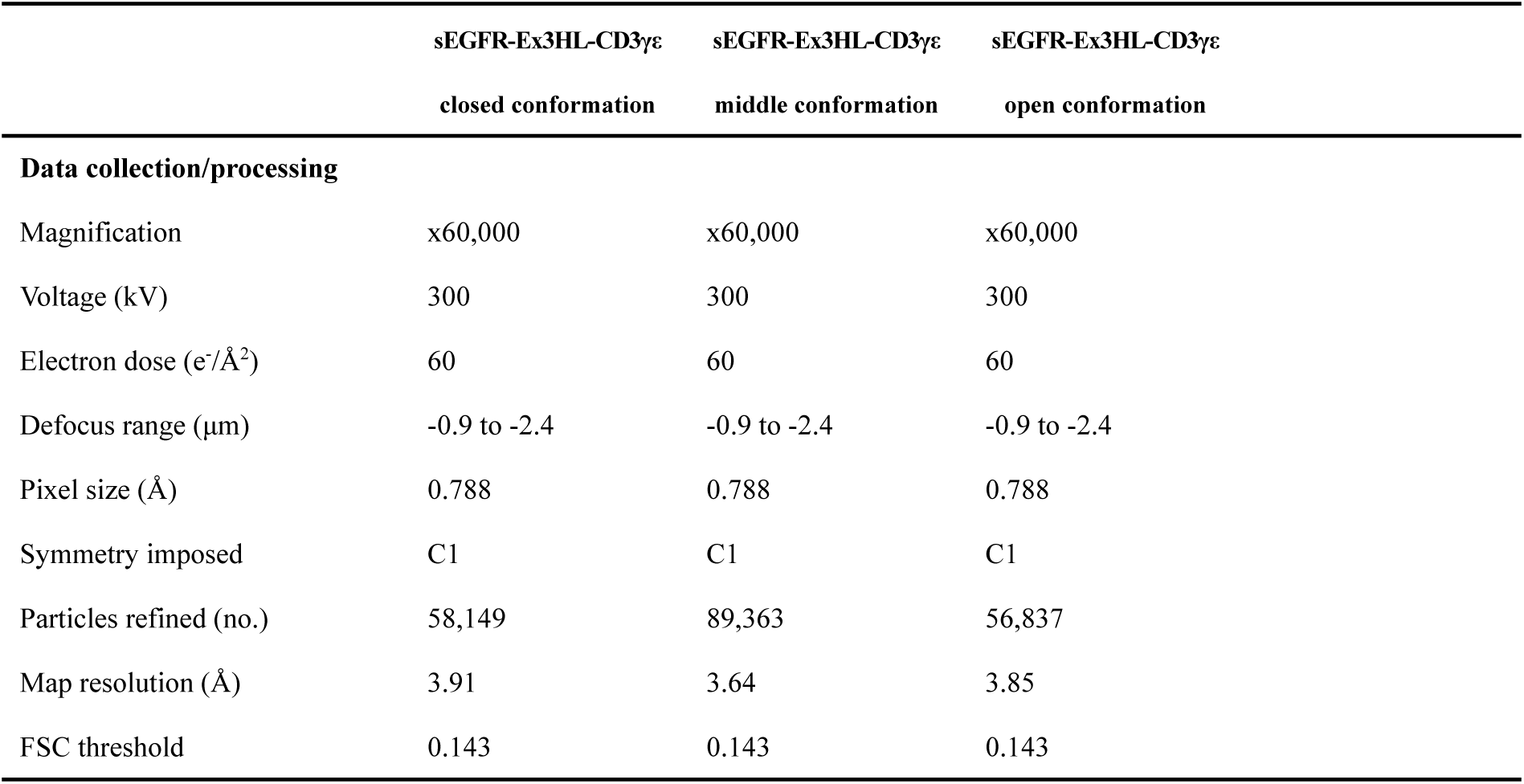
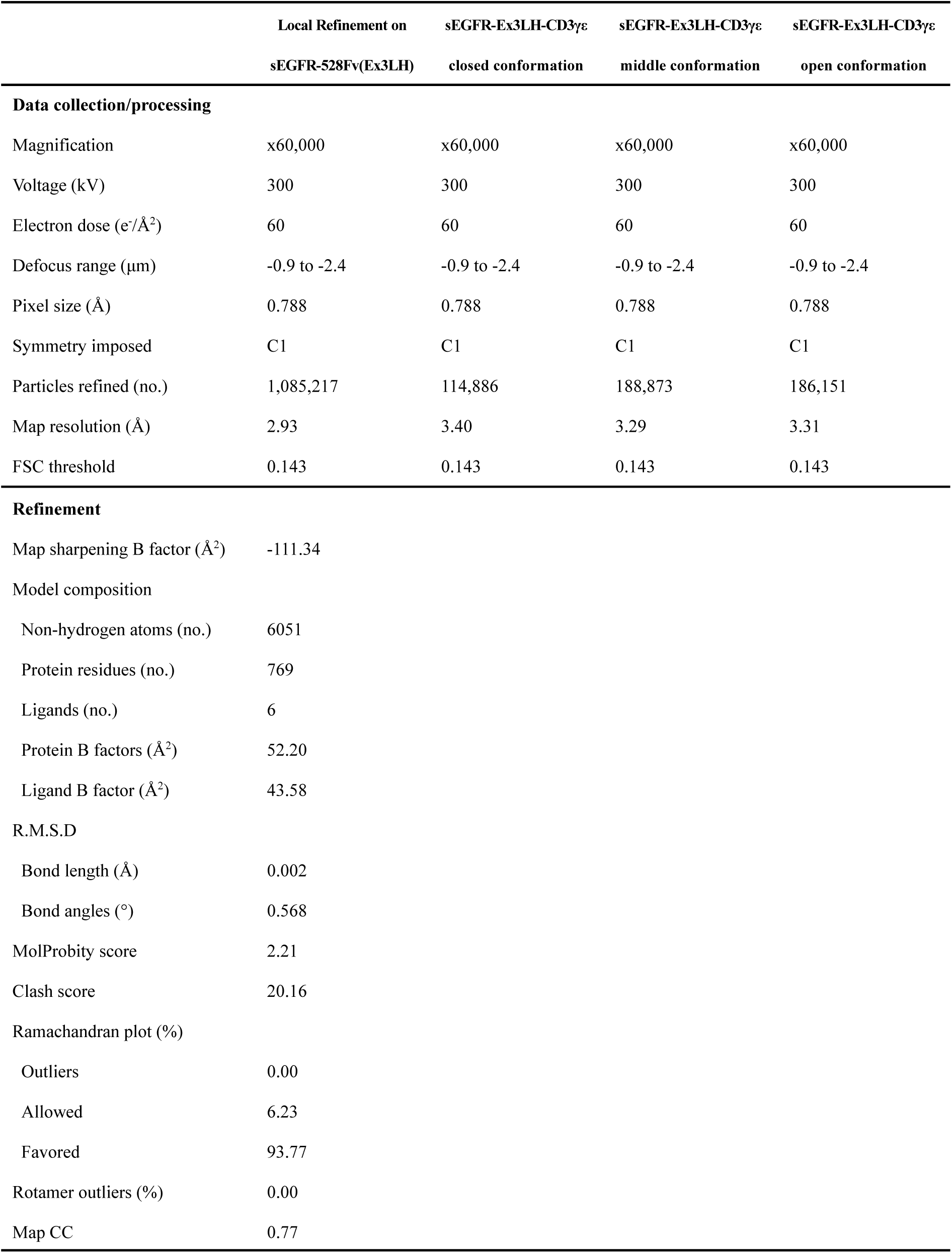
Cryo-EM data collection, refinement, and validation statistics.

**Supplementary Table 3:** Concentration of EGFRD3ND2-mFc and cetuximab Fab for BLI.

|  | Immobilized EGFR<br>concentration (nM) | Binding cetuximab Fab<br>concentration (nM) |
| --- | --- | --- |
| EGFRD3ND2(WT)-mFc | 1000 | 200 |
| S464L-mFc | 475 |  |
| G465R-mFc | 835 |  |
| K467T-mFc | 546 |  |
| S492R-mFc | 493 |  |

**Supplementary Table 4:** Concentration of EGFRD3ND2-mFc and 528 Fab for BLI.

|  | Immobilized EGFR<br>concentration (nM) | Binding 528 Fab<br>concentration (nM) |
| --- | --- | --- |
| EGFRD3ND2(WT)-mFc | 1000 | 125 |
| S464L-mFc | 475 |  |
| G465R-mFc | 835 |  |
| K467T-mFc | 546 |  |
| S492R-mFc | 493 |  |

**Supplementary Video 1: 3DVA of sEGFR-Ex3HL-CD3γε.**

**Supplementary Video 2: 3DVA of sEGFR-Ex3LH-CD3γε.**

